# Evaluation of 21 Brassica microgreens growth and nutritional profile grown under diffrenet red, blue and green LEDs combination

**DOI:** 10.1101/705806

**Authors:** Khaled Y. Kamal, Ahmed A. El-Tantawy, Diaa Abdel Moneim, Asmaa Abdel Salam, Naglaa Qabil, Salwa M. A. I. Ash-shormillesy, Ahmed Attia, Mohamed A. S. Ali, Raúl Herranz, Mohamed A. El-Esawi, Amr A. Nassrallah

## Abstract

Microgreens are rich functional crops with valuable nutritional elements that have health benefits when used as food supplants. Growth characterization, nutritional composition profile of 21 varieties representing 5 species of the *Brassica* genus as microgreens were assessed under light-emitting diodes (LEDs) conditions. Microgreens were grown under four different LEDs ratios (%) (R_80_:B_20_, R_20_:B_80_, R_70_:G_10_:B_20_, and R_20_:G_10_:B_70_). Results indicated that supplemental lighting with green LEDs (R_70_:G_10_:B_20_) enhanced vegetative growth and morphology, while blue LEDs (R_20_:B_80_) increased the mineral composition and vitamins content. Interestingly, combining the nutritional content with the growth yield to define the optimal LEDs setup, we found that the best lighting to promote the microgreen growth was supplying the green LEDs combination (R_70_:G_10_:B_20_). Remarkably, under this proper conditions, Kohlrabi purple, Cabbage red, Broccoli, Kale Tucsan, Komatsuna red, Tatsoi, and Cabbage green had the highest growth and nutritional content profile as microgreens which being a health-promoting in a diet support strategy required for the human health under certain isolated of limited food conditions.

## 1. Introduction

As the world’s population is rapidly growing, with an increasing demand for sustainable sources of food products such as the rich-nutrient functional crops. Ongoing efforts are aimed to find new strategies for food production to meet the demands of the growing world population. Recently, the consumption of microgreens has increased, as a rich-nutrient crop with a high level of nutrition components concentration contains; vitamins, minerals, and antioxidants compared to mature greens, which are helpful in filling the nutritional gap challenges [1]. Furthermore, microgreens being valuable functional crops for their rich-phytonutrients content [2, 3]. Microgreens are a category of edible salad crops that appearing in many upscale markets and restaurants. They are harvested at the base of the hypocotyl when the first true leaves start to emerge, generally, the growth rate is ≤21 days after sowing [4, 5]. Despite their small size, they can provide a high concentration of health-promoting phytochemicals [5]. Commercially greenhouse growers became more interested in the microgreen for their high market levels [4]. Specifically, microgreens of the family *Brassicaceae* have become a popular choice due to its easy way for germination and short growth length and providing wide flavors and colors [5]. Brassicaceae microgreens species could be used as a new ingredient which provides a wide variety of our food [5–7] and valued for containing significant amounts of cancer-fighting glucosinolates [8]. They are also rich in carotenoids, especially lutein, zeaxanthin, and β-carotene [9–11]. Thus, brassica microgreens are considered as a functional food, which serves as a health-promoting or disease preventing supplementals [5, 12]

Several strategies were used and developed for providing optimal greenhouse conditions to increase the microgreen yield. Light emitting diodes (LEDs) is a new light source technology used for greenhouses facilities and space-limited plant growth chambers [13, 14]. It becomes more economically viable with high efficiency and low cost, as well as the ability to select light qualities and intensities [15]. It is reported that crop plants use light for photosynthesis and being responded to the different light intensity, wavelength [16, 17]. Microgreens have a lower demand for photon flux compared to long-cycle crops, thus are ideally adapted to chamber environments. Recently, many studies demonstrated the influence of LEDs (blue or/and red) lighting on the plant vegetative parameters [14, 18, 19] and demonstrated the effect of light quality on the growth of the cultivated plants [8, 20–22]. Nevertheless, a lack of information regarding the combined effect of red and blue and other LEDs lighting such as green light on the plant growth, morphology, and nutrition content profile of microgreens [22, 23]. Furthermore, green light supplies enhance the carotenoid content in mustard microgreens [24].

Although microgreens have been considered as valuable and nutritionally beneficial functional crops, a little is known on the integrity of individual and combined influence of green, red, and blue LEDs on Brassica species microgreens growth and nutritional composition. Therefore, the main purpose of this current study is to define the influence of alternative LEDs light regimens on Brassica species microgreens growth, and nutritional composition and to define which species could serve well as a life support component in many cases. We explore the impact of different four LEDs lighting ratio (Red, Blue, and Green) on 21 Brassica microgreens growth and nutritional profile.

## 2. Material and Methods

### 2.1. Plant Materials and Growth chamber environment

Twenty-one varieties of microgreens representing 5 species of Brassica genus of the *Brassicaceae* family (Table 1) were grown in greenhouse chambers in a collaborated study between the Faculty of Agriculture in both Zagazig University and Cairo University. We used the recommended soil and fertilization properties as reported by [5]. About 10-25 g of seeds, varying based on the seed index of each variety, (Table 1) were sown in peat moss in Rockwool tray in a controlled conditions greenhouse (3 trays per each variety for 3 replicates), cultivated under relative humidity (RH), and carbon dioxide (CO_2_) concentration of 70%, and 500 μmol.mol^−1^, respectively. Each day, 100 ml of CaCl_2_ solution was added to each tray to further stimulate seedling growth. Once cotyledons were fully reflexed 5 d after sowing, 300 ml of 25% nutrient solution was added to each tray daily until harvest. This experiment was carried out simultaneously in the summer season of 2018 from May to September with as a growth length for each species ranging from 6-12 days (Table 1).

**Table 1.** Twenty-one varieties of Brassica microgreens represented 5 species Brassica genera assayed in this study. Growth length (day) and seed index (g) show each variety growth period and the number of seeds used for the sowing.

| Commercial name | Scientific name (genus and species) | Growth length (day) | Seed index (g) |
| --- | --- | --- | --- |
| Broccoli | *Brassica oleracea* L. var. *italica* | 9 | 10 |
| Brussel sprouts | *Brassica oleracea* L. var. *Gemmifera* | 10 | 15 |
| Cabbage green | *Brassica oleracea* L. var. *capitata* f. *alba* | 9 | 10 |
| Cabbage red | *Brassica oleracea* L. var. *capitata* f. *rubra* | 8 | 10 |
| Cabbage savoy | *Brassica oleracea* L. var. *capitata* f. *sabauda* | 8 | 10 |
| Cauliflower | *Brassica oleracea* L. var. *botrytis* | 9 | 15 |
| Collard | *Brassica oleracea* L. var. *viridis* | 10 | 15 |
| Kale Chinese | *Brassica oleracea* L. var. *alboglabra* | 10 | 15 |
| Kale red | *Brassica oleracea* L. var. *acephala* | 9 | 10 |
| Kale Tucsan | *Brassica oleracea* L. var. *acephala* | 9 | 15 |
| Kohlrabi purple | *Brassica oleracea* L. var. *gongylodes* | 7 | 25 |
| Cabbage Chinese | *Brassica rapa* L. var. *pekinensis* | 6 | 15 |
| Komatsuna red | *Brassica rapa* L. var. *perviridis* | 8 | 15 |
| Mizuna | *Brassica rapa* L. var. *nipposinica* | 8 | 15 |
| Pak choy | *Brassica rapa* L. var. *chinensis* | 8 | 15 |
| Rapini | *Brassica rapa* L. var. *ruvo* | 9 | 15 |
| Turnip | *Brassica rapa* L. var. *rapa* | 9 | 10 |
| Mustard Dijon | *Brassica juncea* (L.) Czern. | 12 | 15 |
| Mustard red | *Brassica juncea* (L.) Czern. | 10 | 10 |
| Rutabaga | *Brassica napus* L. var. *napobrassica* | 9 | 10 |
| Tatsoi | *Brassica narinosa* L. var. *rosularis* | 7 | 10 |

### 2.2. LEDs lighting treatments

Brassica microgreen plants were grown under LEDs lighting (Light-emitting diode arrays) were provided by four different light quality ratios (%) treatments of red:blue 80:20 and 20:80 (R_80_:B_20_, and R_20_:B_80_), or red:green:blue 70:10:20, and 20:10:70 (R_70_:G_10_:B_20_, and R_20_:G_10_:B_70_) (Philips GreenPower LED production modules; Koninklijke Philips Electronics, N. V., Amsterdam, The Netherlands), using 0.5 W per LED chip. Each LEDs treatment was carried out in a different room. In the controlled environment greenhouse, the LEDs were placed horizontally, above the bench top, at a height of 50 cm. we adjusted the photosynthetic photon flux density (PPFD) average to 150 μmol.m^−2^. s^−1^ that was provided by the fluorescent lamps and bar-type LEDs. This experiment was performed three times replications with the same conditions.

### 2.3. Harvest, Growth measurements

Microgreen samples were harvested after the growth length for each species (Table1) without seed coats or roots as recommended by [5]. Each replicate used for the measurements consisted of at least 10 grown seedlings. Ten seedlings of each microgreen variety were randomly selected and measured to determine Hypocotyl Length (HL), Leaf Area (LA), for each LEDs treatment. Hypocotyl measurements HL of the harvested seedlings were measured from the tip where the cotyledons split, to the end of the base of the hypocotyl. LA of cotyledons and fully expanded leaves were measured by LA meter (LI-3100; LI-COR Inc. Linclolin, NE) be recording the average of five scans.

Furthermore, another ten randomly selected seedlings for each variety used to assess both, Fresh weight (FW), and Dry weight percentage (DW%). After FW data were measured, samples were oven dried at 80°C for 72 hours. Then DW data were measured. FW and DW values were used to calculate DW% (DW% = (DW/FW × 100).

### 2.4. Elemental Analysis

Fresh microgreens (50 g FW per each sample) were collected and rinsed 3X using H2Odd to remove any surface residue. Dried microgreens (2◻g per replicate) were grounded into a fine powder to analysis the elemental composition., Each of the 21 samples was subjected to acid digestion procedures and quantitative measurements of the following elements: P, K, Ca, Na, Fe, Mn, Cu, and Zn were done using inductively coupled plasma optical emission spectrometry (ICP-OES) following the methods of Huang and Schulte [25]. To assure the accuracy of the method, standard reference materials (Apple leaves, NIST® SRM® 1515, NIST1515, SIGMA, USA, and Spinach leaves, NIST® SRM® 1570a, NIST1570A) were used and evaluated using the same digested procedure. For each ICP-OES analyte, the limit of detection (LOD) and limit of quantification (LOQ), which are a function of the sample mass were determined (Supplementary Table 1)

### 2.5. Vitamin and Carotenoid concentration analysis

#### 2.5.1. Phylloquinone

Phylloquinone was determined according to a previously reported method by [26]. Under dime light, 0.2 g of dried microgreens were homogenized in 10 mL of H2O and 0.4 mL of 200 μg/mL menaquinone used as an internal standard. The sample was supplied with 15 mL of 2-propanol/hexane (3:2 v/v) and were vortexed for 1 min. Then the sample was centrifuged at 1500g at 21°C for 5 min. Then we transferred the upper layer (hexane) into a new glass tube and to dry using a stream of N2. The residues of the sample were dissolved using 4 mL of hexane. Then, to purify the extract, 1 mL of the dissolved extract was loaded onto preconditioned silica gel columns (4 mL of 3.5% ethyl ether in hexane, followed by 4 mL of 100% hexane). We used 2 mL of hexane to wash the columns. Phylloquinone was eluted with 8 mL of 3.5% ethyl ether in hexane and then evaporated at 40 °C under N2 flow. Further, it is reconstituted in 2 mL of mobile phase (99% methanol and 1% 0.05 M sodium acetate buffer, pH = 3.0) and is filtered through a 0.22 μm nylon syringe filter (Millipore, Bedford, MA). To detect the phylloquinone, we used a photodiode array detector (DAD) (G1315C, Agilent, Santa Clara, CA) on Agilent 1200 series HPLC system and absorbance wavelength was 270 nm. 20 μL of the extract was injected into the HPLC and being run through a C18 column (201TP, 5 μm, 150 × 4.6 mm, Grace, Deerfield, IL) flowing at the rate of 1 mL/ min. The phylloquinone content was measured according to the internal standard method based on peak areas.

#### 2.5.2. Carotenoids and Tocopherols

To extract both carotenoids and tocopherols, we followed the procedure described by [27] and modified by Xiao et al. [5]. In 15 mL screw-cap glass vial, 0.05 g of dried fine powder was homogenized in 7.5 mL of 1% butylated hydroxytoluene in ethanol and 500 μL of 86.82 μM trans-βapo-8 carotenal as an internal standard was added. 180 μL of 80% KOH was supplied to the mixture and, the vials were capped and placed in a dry bat at 70 °C for 15 min. The vials were removed and being cooled on ice 4°C for 5 min. The mixture was transferred into 15 mL centrifuge tubes supplied with 3.0 mL of deionized water and 3.0 mL of hexane/toluene solution (10:8 v/v). The mixture was carefully vortexed for 1 min and then were centrifuged at 1000g for 5 min. After centrifugation, the upper organic layer was collected into an 8 mL glass culture tube and immediately placed into a nitrogen evaporator set at 30 °C. on the other hand, the lower layer was extracted with 3.0 mL of hexane/toluene (10:8 v/v). this extraction process was repeated at least four times until the upper layer is colorless. After evaporation, the residue was diluted in 500 μL of mobile phase acetonitrile/ethanol (1:1 v/v), filtered into an HPLC amber vial using nylon filter (0.22 μm, Millipore, Bedford, MA). Subsequently, 20 μL was inoculated for HPLC analysis. Carotenoid and tocopherol concentrations were measured using isocratic reverse-phase high-performance liquid chromatography (RP-HPLC). Absorbance was measured at 290 nm (for tocopherols) and 450 nm (for carotenoids).

#### 2.5.3. Ascorbic Acid

Total ascorbic acid (TAA) was assessed spectrophotometrically according to [28]. 3g fresh microgreens were homogenization in 10 mL of ice-cold 5% metaphosphoric acid (w/v) at 4°C at 15000 rpm for 1 min. The homogenized then centrifuged at 7000g for 20 min at 4°C. The supernatant was filtered through Whatman 4# filter paper. TAA was measured spectrophotometrically at 525 nm. Concentrations of TAA was calculated from an L-ascorbic acid standard curve.

### 2.6. Clustering hierarchical analysis

In order to extrapolate the similarities and the dissimilarities among the 21 microgreens in growth and nutritional assessment, hierarchical cluster analysis was performed using the normalized data set using *class Orange clustering hierarchical* using *ORANGE version 3.7* [29].

### 2.7. Statistical analysis

The experiment was laid out in a randomized block design in a factorial arrangement with LEDs (four levels) and Microgreens (Twenty-one varieties) for three different biological replicates. Data were collected and analyzed according to [30]. SPSS_*v.22*_ software was used to analyze the variance of differences using ANOVA test statistically followed by LSD analysis. The degree of freedom was followed as *P*≤0.05, *P*≤0.01, and *P*≤0.001 considers the statistical significance and represents as *, **, *** respectively.

## 3. Results

### 3.1. The influence of LEDs on microgreens growth, nutritional profile

In the present work, four different LEDs lighting ratio (%) treatments of R_80_:B_20_, R_20_:B_80_, R_70_:G_10_:B_20_, and R_20_:G_10_:B_70_ were implemented. Growth parameters of the 21 varieties were analyzed (Fig. 1). Results revealed that those microgreens are grown under the R_70_:G_10_:B_20_ had the highest growth and morphology targeted parameter, while the lowest growth parameters were observed under R_20_:B_80_ (Figure 1). The Hypocotyl length (HL) and leaf area (LA) of the microgreens were significantly elongated in plants grown under R_70_:G_10_:B_20_ compared to those grown under R_80_:B_20_, R_20_:B_80_, and R_20_:G_10_:B_70_, respectively (Figure1)., Fresh weight (FW) and Dry weight % (DW%) of those microgreens grown under R_70_:G_10_:B_20_ treatment showed the highest values; on average; 0.4g (FW), and 6.27 %(DW%). Indicating that R_70_:G_10_:B_20_ combination induces an increase in all studied growth and morphology parameters in comparison with the other LEDs lighting treatments (Figure1).

**Figure 1.**
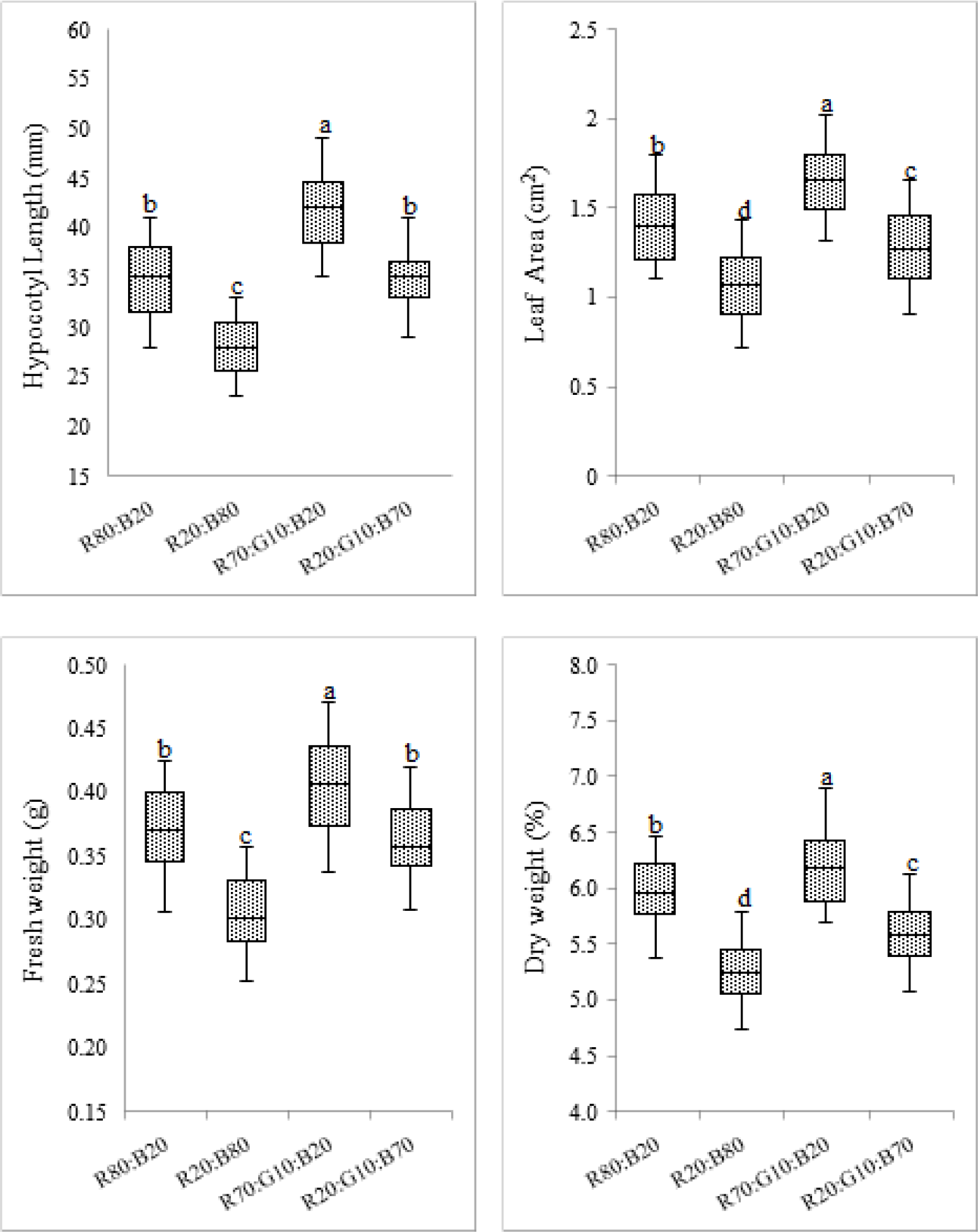
Box plot of growth and morphological measurements of *Brassica* microgreens grown under LEDs treatments. The plot illustrates the Mean and median of (Hypocotyl Length (mm), Leaf Area (cm^2^), Fresh weight (g), and Dry weight (%)) of 21 varieties of Brassica microgreens represented 5 species grown under different light-emitting diodes (LEDs) ratio (%) of red:blue 80:20 (R_80_:B_20_), red:blue 20:80 (R_20_:B_80_), red:green:blue 70:10:20 (R_70_:G_10_:B_20_), or red:green:blue 20:10:20 (R_20_:G_10_:B_70_) (Supplemental Table 2 and 3). Resulting ranking could be analyzed with point values of Mean and Median or uncertainty range with box. Statistical analysis is performed using a one-way ANOVA test (P ≤ 0.05). Small letters denote statistically significant differences.

Considering that R_70_:G_10_:B_20_ LEDs lighting combination has an impact in targeted growth parameters, we investigated whether it has a functional influence on the nutritional composition profile by conducting an ICP analysis of macro and microelements from 21 varieties Brassicaceae microgreen using lowest growth enhancer combination as internal references. Unexpectedly, relative macro and microelements content were showed a dramatic decreased compared to R_20_:B_80_ and the other LEDs ratios (Figure 2 and 3). While the highest levels were obtained in microgreens were growing under R_20_:B_80_ combination. However, the analysis also did not take the yield into consideration.

**Table 2.** Growth, and nutritional composition profile of highest Brassica microgreens grown under light-emitting diodes (LEDs) ratio (%) of red:green:blue 70:10:20 (R_70_:G_10_:B_20_). List of the 7 brassica microgreens is exported from the hierarchical cluster analysis (Figure 7).

|  | **Kohlrabi**<br>**purple** | **Cabbage**<br>**red** | **Broccoli** | **Kale**<br>**Tucson** | **Komatsuna**<br>**red** | **Tatsoi** | **Cabbage**<br>**green** |
| --- | --- | --- | --- | --- | --- | --- | --- |
| Hypocotyl Length (mm) | 46 | 47 | 42 | 49 | 44 | 46 | 45 |
| Leaf Area (cm<sup>2</sup>) | 1.82 | 1.93 | 1.65 | 2.02 | 1.75 | 1.86 | 1.83 |
| Fresh weight (g) | 0.44 | 0.46 | 0.42 | 0.47 | 0.41 | 0.45 | 0.44 |
| Dry weight (%) | 6.65 | 6.18 | 6.55 | 6.90 | 6.57 | 6.34 | 6.25 |
| P (mg/100 g FW) | 68 | 62 | 59 | 63 | 66 | 64 | 62 |
| K (mg/100 g FW) | 322 | 224 | 319 | 280 | 320 | 300 | 183 |
| Ca (mg/100 g FW) | 65 | 84 | 92 | 55 | 53 | 44 | 87 |
| Na (mg/100 g FW) | 46 | 40 | 50 | 46 | 28 | 35 | 69 |
| Fe (mg/100 g FW) | 0.77 | 0.69 | 0.74 | 0.76 | 0.76 | 0.65 | 0.67 |
| Zn (mg/100 g FW) | 0.43 | 0.40 | 0.42 | 0.38 | 0.38 | 0.41 | 0.33 |
| Cu (mg/100 g FW) | 0.08 | 0.10 | 0.11 | 0.07 | 0.05 | 0.08 | 0.06 |
| Mn (mg/100 g FW) | 0.39 | 0.35 | 0.41 | 0.46 | 0.34 | 0.35 | 0.37 |
| Phylloquinone<br>(ug/ g FW) | 2.6 | 2.3 | 2.0 | 1.7 | 2.2 | 1.5 | 1.4 |
| α-tocopherol<br>(mg/100 g FW) | 17.6 | 29.5 | 17.3 | 19.4 | 25.0 | 29.3 | 14.3 |
| Total Ascorbic Acid<br>(mg/100 g FW) | 77.1 | 127.4 | 84.6 | 73.2 | 97.5 | 99.5 | 118.9 |
| β-carotene<br>(mg/100 g FW) | 6.6 | 9.9 | 6.9 | 5.4 | 7.1 | 10.6 | 9.6 |

**Figure 2.**
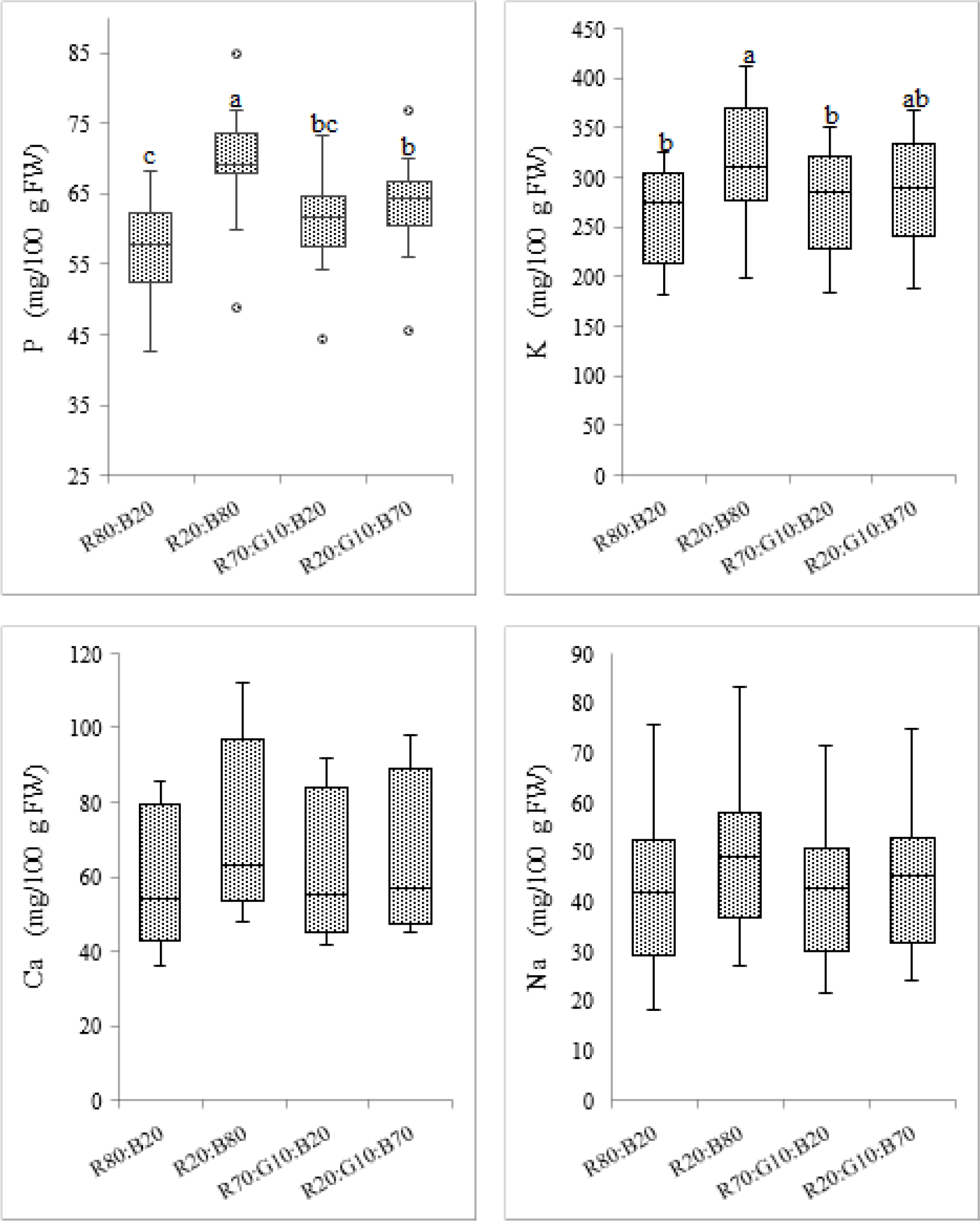
Box plot of mineral composition and content of macroelements of *Brassica* microgreens grown under LEDs treatments. The plot illustrates the Mean and median of macroelements concentrations; P, K, Ca and Na (mg/100 g FW) of 21 varieties of Brassica microgreens represented 5 species grown under different light-emitting diodes (LEDs) ratio (%) of red:blue 80:20 (R_80_:B_20_), red:blue 20:80 (R_20_:B_80_), red:green:blue 70:10:20 (R_70_:G_10_:B_20_), or red:green:blue 20:10:20 (R_20_:G_10_:B_70_) (Supplemental Table 4 and 5). Resulting ranking could be analyzed with point values of Mean and Median or uncertainty range with box. Statistical analysis is performed using a one-way ANOVA test (P ≤ 0.05). Small letters denote statistically significant differences.

**Figure 3.**
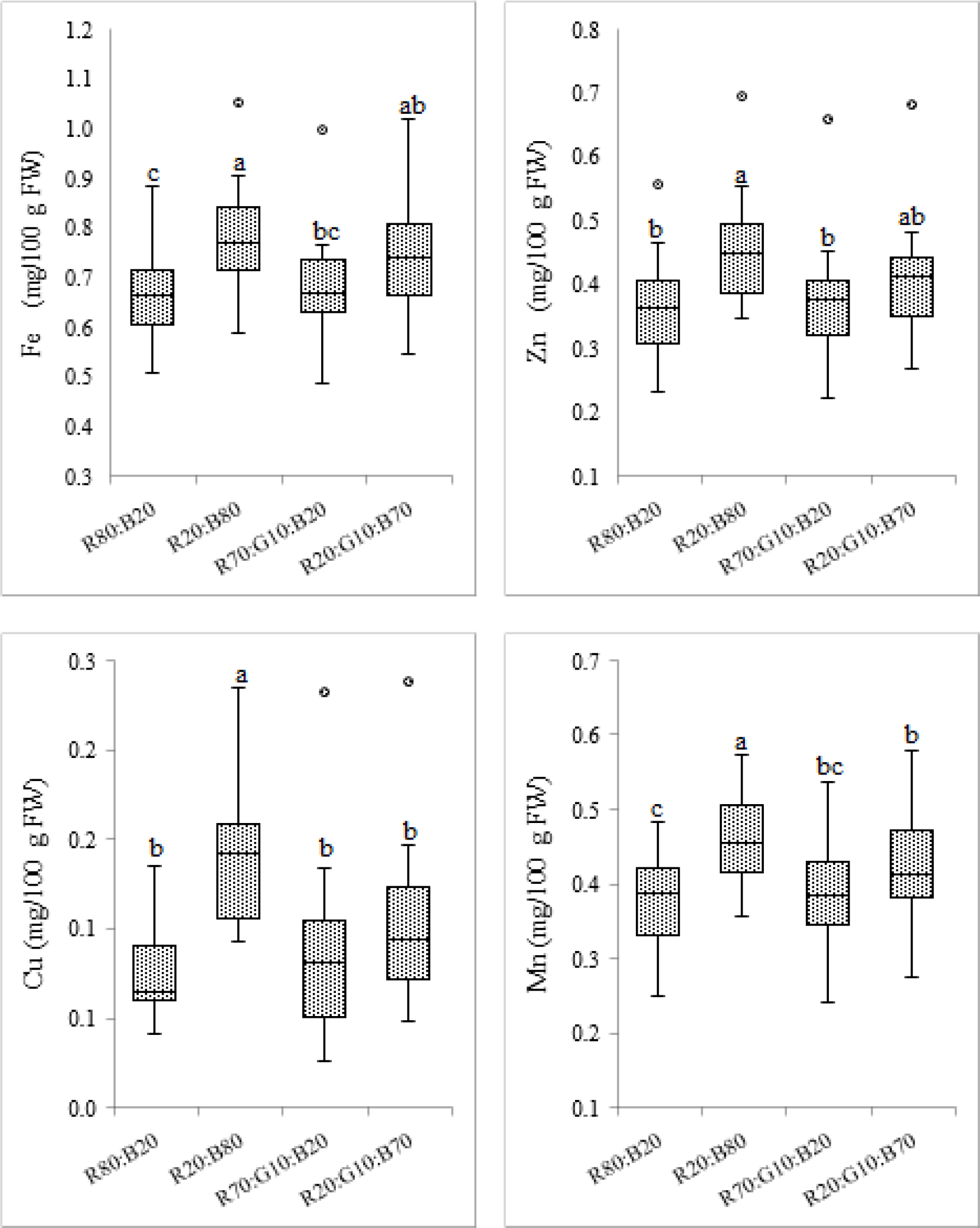
Box plot of mineral composition and content of microelements of *Brassica* microgreens grown under LEDs treatments. The plot illustrates the Mean and median of microelements concentrations; Fe, Zn, Cu and Mn (mg/100 g FW) of 21 varieties of Brassica microgreens represented 5 species grown under different light-emitting diodes (LEDs) ratio (%) of red:blue 80:20 (R_80_:B_20_), red:blue 20:80 (R_20_:B_80_), red:green:blue 70:10:20 (R_70_:G_10_:B_20_), or red:green:blue 20:10:20 (R_20_:G_10_:B_70_) (Supplemental Table 6 and 7). Resulting ranking could be analyzed with point values of Mean and Median or uncertainty range with box. Statistical analysis is performed using a one-way ANOVA test (P ≤ 0.05). Small letters denote statistically significant differences.

Considering the influence of LEDs lighting combination on the microgreen’s growth together with nutrition components value, we analyzed deeper the vitamin and carotenoid contents. Agreeing with our previous result obtained in the macro- and microelements, we found that the contents of Phylloquinone, α-tocopherol, Total Ascorbic Acid (TAA), and β-carotene of 21 varieties of Brassica microgreens grown under red: blue 80:20 (R_80_:B_20_), were significantly increased compared to other combination respectively (Figure 4).

**Figure 4.**
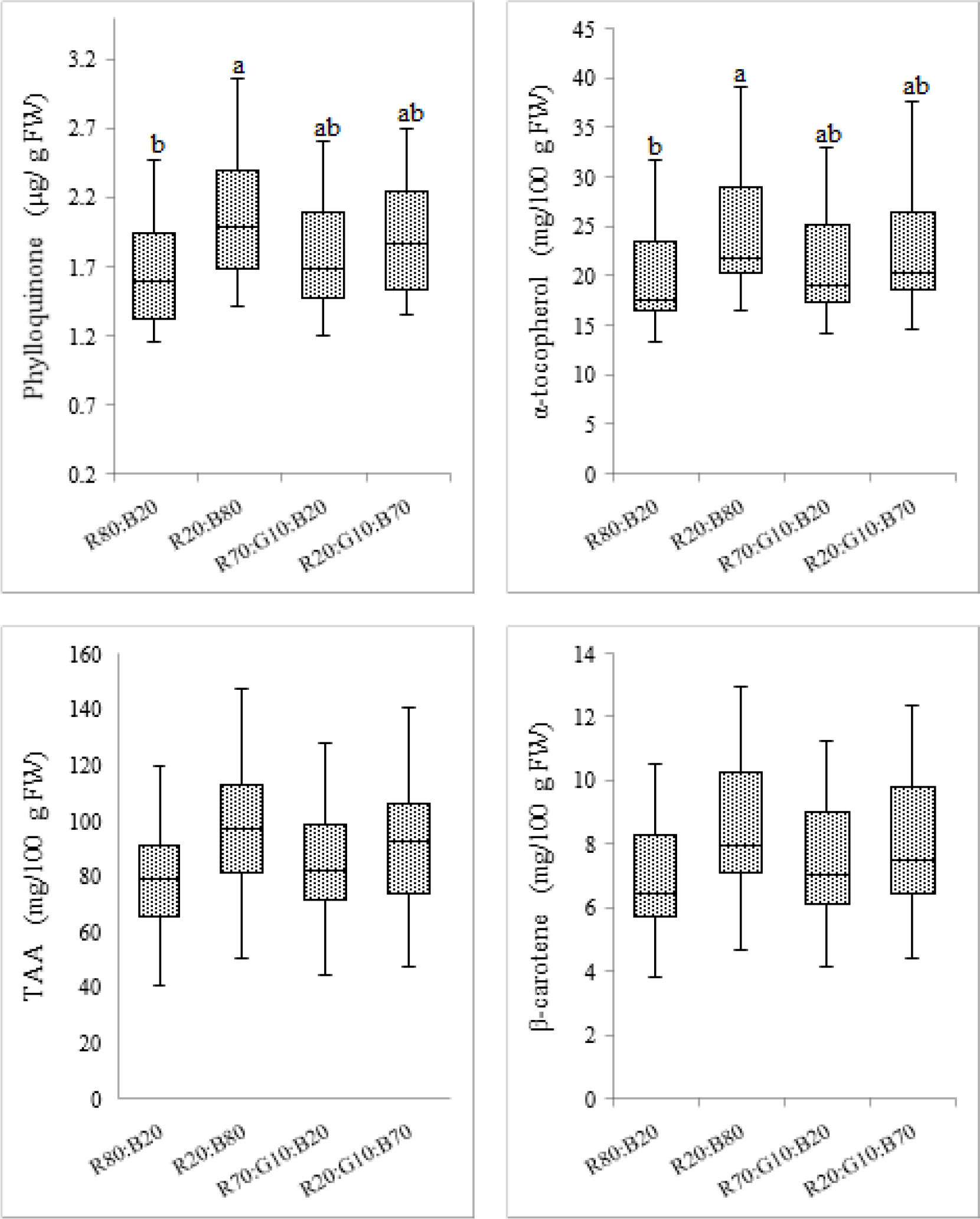
Box plot of vitamin and carotenoid concentrations of *Brassica* microgreens grown under LEDs treatments. The plot illustrates the Mean and median of vitamin and carotenoids concentrations; Phylloquinone (ug/ g FW), α-tocopherol, Total Ascorbic Acid (TAA), and β-carotene (mg/100 g FW) of 21 varieties of Brassica microgreens represented 5 species grown under different light-emitting diodes (LEDs) ratio (%) of red:blue 80:20 (R_80_:B_20_), red:blue 20:80 (R_20_:B_80_), red:green:blue 70:10:20 (R_70_:G_10_:B_20_), or red:green:blue 20:10:20 (R_20_:G_10_:B_70_) (Supplemental Table 8 and 9). Resulting ranking could be analyzed with point values of Mean and Median or uncertainty range with box. Statistical analysis is performed using a one-way ANOVA test (P ≤ 0.05). Small letters denote statistically significant differences.

### 3.2. Conclude the optimum LEDs conditions for Brassica microgreens growth conditions

Our previous data showed that LEDs lighting combination has an impact on all growth and nutritional parameters. More precisely. We found that Brassica microgreen varieties were grown under the LEDs lighting of R_70_:G_10_:B_20_ combination enhances the Hypocotyl length, leaf area, fresh weight, and dry weight compared to other LEDs combination. While minerals (macro and microelements) and vitamins were significantly increased corresponding to plants grown under R_80_:B_20_. Attempts to detect the best LEDs combination taking into consideration the actual yield of microgreens, we conducted a correlation analysis with the yield. We estimated the minerals and vitamins concentrations corresponding to the actual fresh weight yielded (Figure 5 and 6). Interestingly, we found that mineral compositions and vitamins content in the yielded fresh weight were significantly increased in the microgreen varieties grown under the LEDs lighting of R_70_:G_10_:B_20_ combination compared to other combination (Figure 5 and 6).

**Figure 5.**
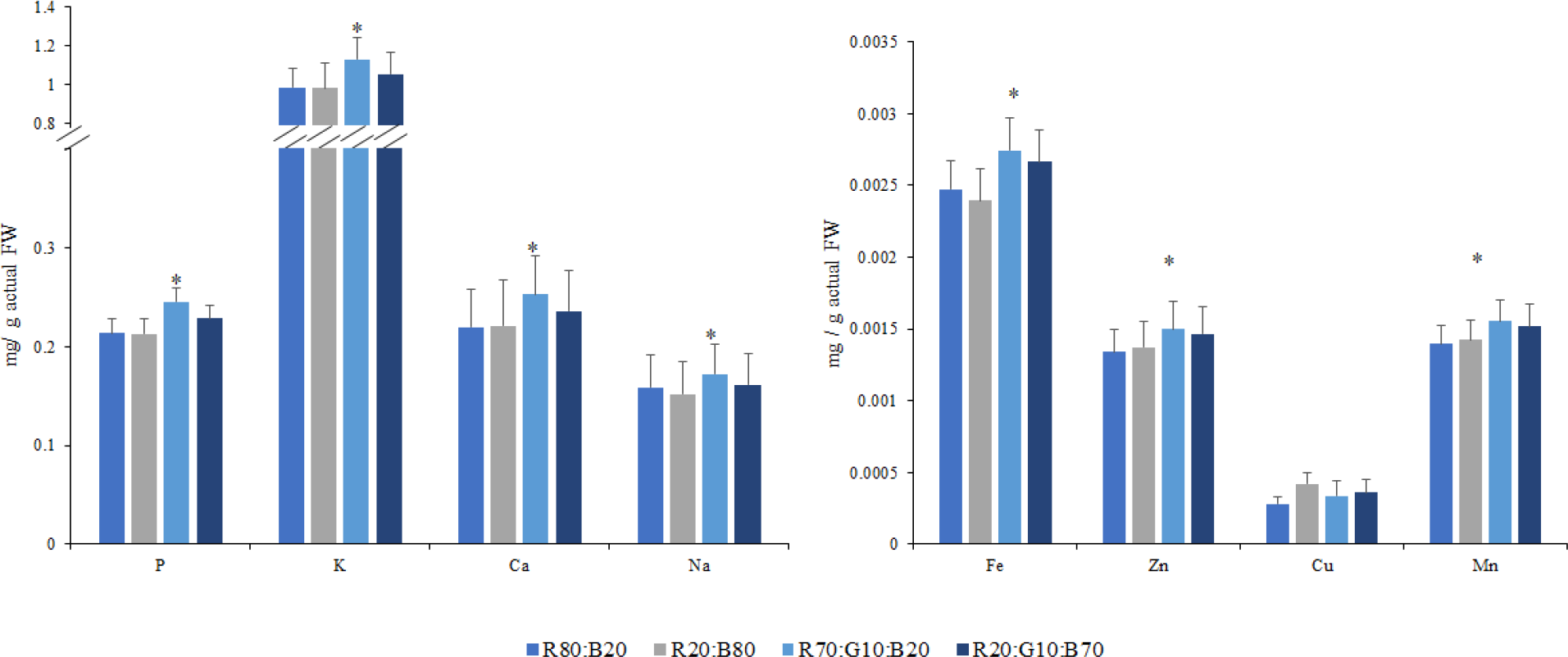
Mineral composition and content of *Brassica* microgreens under LEDs treatments. **A)** Mean macroelement concentration of P, K, Ca, and Na. **B)** Mean microelement concentration of Fe, Zn, Cu, and Mn of 21 verities of Brassica microgreens exposed to different light-emitting diodes (LEDs) ratio (%) of red:blue 80:20 (R_80_:B_20_), red:blue 20:80 (R_20_:B_80_), red:green:blue 70:10:20 (R_70_:G_10_:B_20_), or red:green:blue 20:10:20 (R_20_:G_10_:B_70_). Data represents as a mean concentration corresponding to the actual Fresh weight (fresh weight results of each LEDs treatments of the 21 verities (Supplementary Table 3) (Actual concentration (mg/ g actual FW) = Concentration (mg/ 100 g FW) X Fresh weight (g) / 100). Mean±SE values are based on a representative sample from each treatment across three experimental replications. * for significant at P ≤ 0.05.

**Figure 6.**
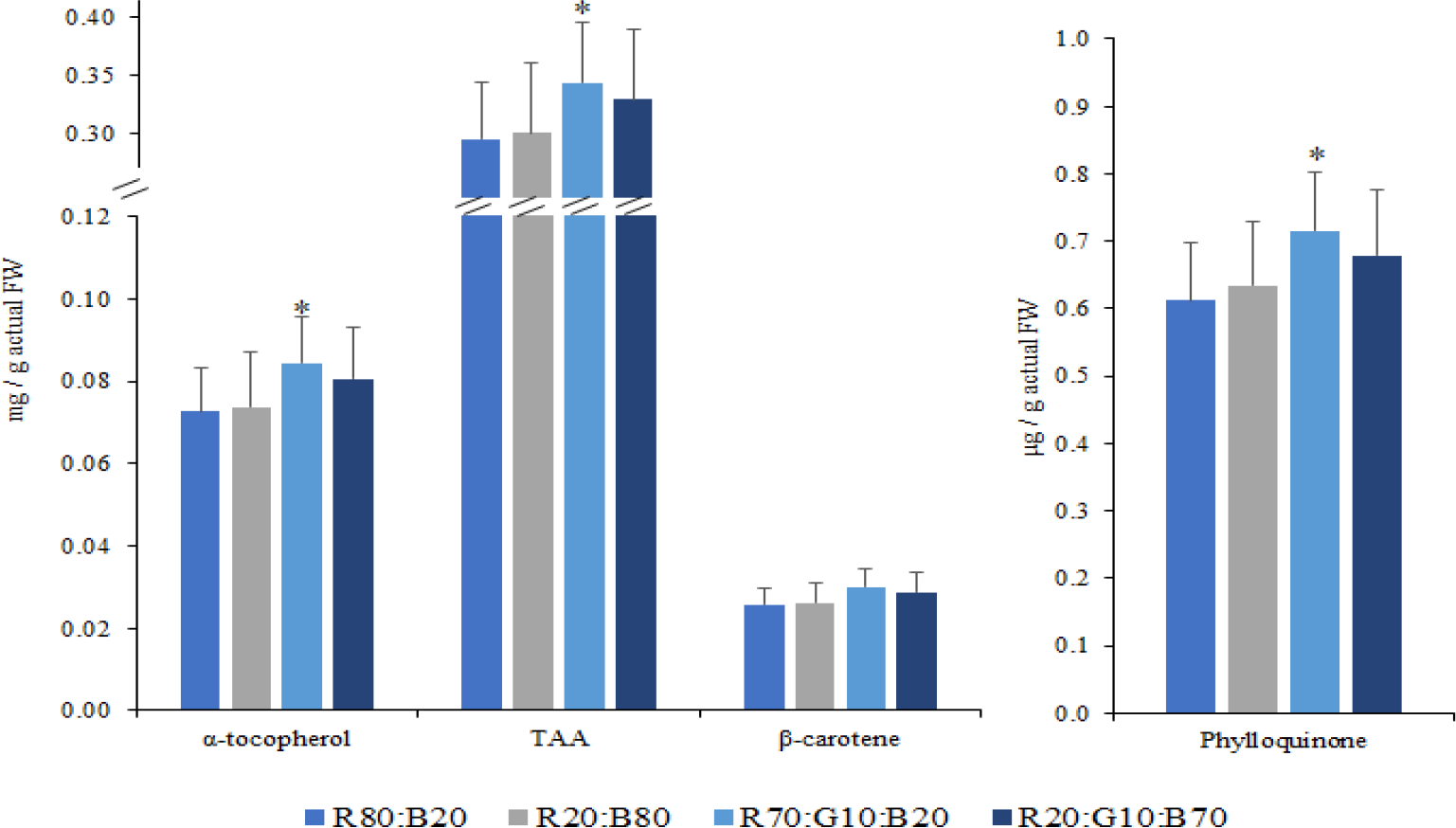
Assessment of vitamin and carotenoid concentrations of *Brassica* microgreens under LEDs treatments. **A)** Mean α-tocopherol, Total Ascorbic Acid (TAA), and β-carotene (mg/100 g FW) concentration. **B)** Mean Phylloquinone (ug/ g FW) concentration of 21 verities of Brassica microgreens exposed to different light-emitting diodes (LEDs) ratio (%) of red:blue 80:20 (R_80_:B_20_), red:blue 20:80 (R_20_:B_80_), red:green:blue 70:10:20 (R_70_:G_10_:B_20_), or red:green:blue 20:10:20 (R_20_:G_10_:B_70_). Data represents as a mean concentration corresponding to the actual Fresh weight (fresh weight results of each LEDs treatments of the 21 verities (supplementary Table 3) (Actual concentration (mg/ g actual FW) = Concentration (mg/ 100 g FW) X Fresh weight (g) / 100). For Phylloquinone ((Actual concentration (mg / g actual FW) = Concentration (μg/ g FW) X Fresh weight (g)). Mean±SE values are based on a representative sample from each treatment across three experimental replications. * for significant at P ≤ 0.05.

### 3.3. Hierarchical cluster analysis of 21 varieties of Brassica microgreens

A hierarchical cluster analysis profiled growth, mineral compositions and vitamins content of 21 microgreens varieties grown under R_70_:G_10_:B_20_ family has been performed using *class Orange clustering hierarchical* using *ORANGE version 3.7* [29]. Presented data of microgreens grown under R_70_:G_10_:B_20_, which present the highest values of growth and nutritional profile are shown in Figure 7 and Table 2. We utilized the hierarchical analysis methods (average-linkage distance between two clusters) to evaluate whether these trends were consistent across the 21 varieties under study. The hierarchical cluster analysis shows that the 21 microgreens are classified into five groups. Among the five groups, the highest distance group (Figure 7, cluster group in yellow color + Kale Tucsan in green cluster) contained 7 microgreens(Kohlrabi purple, Cabbage red, Broccoli, Kale Tucsan, Komatsuna red, Tatsoi, Cabbage green) which are representing 3 species (*B. oleracea, B. rapa, B. narinosa*).

**Figure 7.**
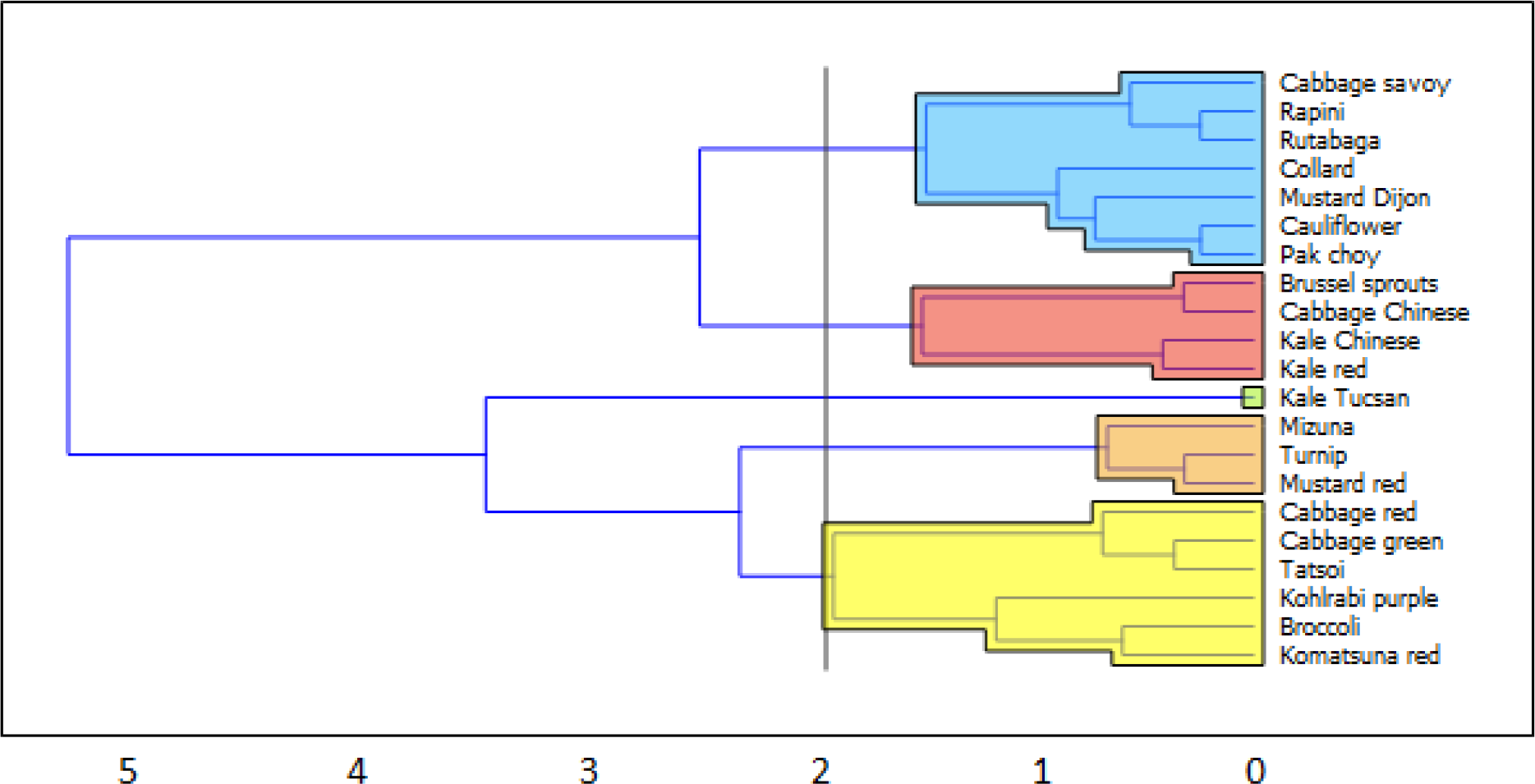
The average-linkage on the normalized data sets of mineral composition and vitamin and carotenoid concentrations corresponding to the actual fresh weight by means of the Hierarchical method using growth and morphology measurements data of 21 varieties Brassica microgreens grown under light-emitting diodes (LEDs) ratio (%) of red:green:blue 70:10:20 (R_70_:G_10_:B_20_). The complete profile of the highest cluster value (Yellow cluster) microgreens presented in Table 2.

## 4. Discussion

Due to the increased interest with providing the controlled environment greenhouses with LEDs lighting and for increasing the microgreen growth and nutritional profile, we investigated the impact of four different LEDs lighting ratios on the growth and nutritional quality assessment of 21 varieties belong to Brassica genera of the family Brassicaceae grew as microgreens. Microgreens are reported in many studies as valuable source vitamins, phenolics and mineral compositions [31]. Enhancing their nutritional qualities and growth is an exciting avenue of research and agriculture biotechnology.

In our study, we reveal various effects on the combination ratios between blue LEDs, red and green LEDs. A plant grown under a monochromatic light beam also stimulate specific photoreceptors that are involved in numerous biological processes. Enhance the nutritional profile and plant growth was demonstrated in many species, such as rice [32], Brassica spp. [5, 17, 22], etc. It has been reported that red and blue light are important for the expansion of the hypocotyl elongation, pigments accumulation, and enhancement of biomass [33]. In contrast, exposure to green LEDs increases biomass at a high intensity [34]. We notice that growing microgreens under R_70_:G_10_:B_20_ shows the highest value of the vegetative parameters, taking in our consideration the yield produced under all combination treatments (Figure 1). These results provide a clear indication that blue LEDs in combination with red LEDs and high-intensity green LEDs are more efficient for plants microgreen growth. Providing green lighting within the growing conditions enhance Brassica microgreens growth, while, increasing the blue light ratio had a passive response to the growth.

Many reports demonstrate the positive influence of the red:blue lighting plant growth and photosynthesis [13, 16, 17, 21, 24, 35, 36]. Furthermore, a red, blue, and green light combination has an effective source for photosynthesis [37].

Consequently, supply the red and blue LEDs combination with a green light has a significant impact on lettuce leaves growth and photosynthetic rate compared with the red and blue LEDs only. [38, 39]. It appears that blue and red light enhances the anthocyanins accumulation in leaves and become black, while green light stimulates phytochrome, shifting the active pool of Type I and Type II phytochromes to include reverse accumulation of anthocyanins [40].

Consequently, we demonstrate the positive influence of providing green light improving microgreens growth and morphology. It is reported that HL and LA of kohlrabi, mizuna, and mustard were increased when grown under green light R_74_:G_18_:B_8_ compared with the R_87_:B_13_, while FW and DW were greater of those microgreens grown under providing green light than blue/red [41]. Moreover, FW of broccoli microgreens grown under light ratios of R_85_:G_10_:B_5_ and R_80_:B_20_ were significantly increased than under R_70_:G_10_:B_20_ [42]. The same influence is observed on chlorophyll content which improved significantly of the plant grown under additional green light [22, 41] Furthermore, the reduction on the growth parameters due to the increased of blue lighting was reported. It is found that the hypocotyl elongation of kohrabi, mizuna, mustard was decreased under the red:blue light combination due to the inhibition of the gibberellins (GA), which inhibit the hypocotyl elongation [36].

Growing microgreens plants under blue LEDs R_20_:B_80_ in our study enhance the minerals composition and the vitamin content accompanied with the less growth yield compared with the green LEDs R_70_:G_10_:B_20_. It is reported that broccoli microgreens grown under blue light (R_0_:B_100_) produce higher nutrient-dense microgreens [8, 42]. Blue light could be shown as a dominant means of regulating the nutrient content synthesis such as proton pumping, ion channel, activities, and membrane permeability [41, 42].

Comparing the LEDs lighting ratios to conclude the proper conditions, we accompanied the nutritional profile with the actual growth yielding. We found that green LEDs R_70_:G_10_:B_20_ has the proper yielded influence and produced final higher mineral concentration and vitamin content due to the high growth yield. Despite the blue LED treatment to increase the mineral and vitamin content, but it is accompanied by less growth yield.

In conclusion, the assessment of 21 brassica microgreens growth and nutritional profile grown under LEDs technology provides a satisfactory growing conditions examination of microgreens. Providing green lighting ratio of R_70_:G_10_:B_20_ show a positive influence on the growing microgreens growth and morphology. Interestingly, Kohlrabi purple, Cabbage red, Broccoli, Kale Tucsan, Komatsuna red, Tatsoi, Cabbage green are presented as the top microgreen’s candidates of our study assessment that serve as a life support component in limited space-based conditions and controlled environment greenhouse.

## Acknowledgments

Special thanks to DAAD and Exceed Swindon to provide this opportunity to present this work in the EXCEED SWINDON EXPERT WORKSHOP (Aswan-EGYPT 2018). Furthermore, Many thanks to all the lab member in the Agronomy Department, faculty of agriculture, Zagazig university for their technical support.

## Author contributions

K.Y.K., A.A.N conceived and designed the experiments. K.Y.K., A.A.E, A.A.N, M.A.S.A. and S. J. L.Z. performed the experiment. K.Y.K., D.A.M., A.A.S, N.Q., S.M.A and S.Y.M analyzed the data. K.Y.K., A.A.N. wrote the manuscript. R.H., M.A.E. contributed to the manuscript writing and revision. All authors revised the manuscript.

