## Supplemental data for "Evaluation of 21 Brassica microgreens growth and nutritional profile grown under diffrenet red, blue and green LEDs combination"

**Supplemental Table 1.** Limit of detection (LOD/mg/kg DW) and limit of quantification (LOQ/mg/kg DW) based on background equivalent concentration.

| | LOD mg/Kg DW | LOQ mg/Kg DW |
|---|---|---|
| P | 0.0018 | 0.005 |
| K | 25.0068 | 80.004 |
| Ca | 0.0108 | 0.055 |
| Mg | 0.0004 | 0.016 |
| Na | 0.0116 | 0.204 |
| Fe | 0.0034 | 0.002 |
| Zn | 0.0004 | 0.006 |
| Cu | 0.0002 | 0.006 |

**Supplemental Table 2.** Growth and morphological measurements (Hypocotyl Length (mm) and Leaf Area (cm2)) of 21 varieties of Brassica microgreens represented 5 species grown under different light-emitting diodes (LEDs) ratio (%) of red:blue 80:20 ($R_{80}$:$B_{20}$), red:blue 20:80 ($R_{20}$:$B_{80}$), red:green:blue 70:10:20 ($R_{70}$:$G_{10}$:$B_{20}$), or red:green:blue 20:10:20 ($R_{20}$:$G_{10}$:$B_{70}$). Mean±SE values are based on a representative sample from each treatment across three experimental replications. NS, *, **, ***Nonsignificant or significant at P ≤ 0.05, 0.01, or 0.001, respectively.

| Commercial name | Hypocotyl Length (mm) | | | | Leaf Area ($cm^2$) | | | |
|---|---|---|---|---|---|---|---|---|
| | $R_{80}$:$B_{20}$ | $R_{20}$:$B_{80}$ | $R_{70}$:$G_{10}$:$B_{20}$ | $R_{20}$:$G_{10}$:$B_{70}$ | $R_{80}$:$B_{20}$ | $R_{20}$:$B_{80}$ | $R_{70}$:$G_{10}$:$B_{20}$ | $R_{20}$:$G_{10}$:$B_{70}$ |
| Broccoli | 37.4 | 27.2 | 42.2 | 33.1 | 1.42 | 1.07 | 1.65 | 1.26 |
| Brussel sprouts | 34.2 | 27.6 | 39.6 | 36.3 | 1.29 | 0.94 | 1.52 | 1.13 |
| Cabbage green | 38.5 | 29.7 | 45.8 | 37.3 | 1.63 | 1.25 | 1.83 | 1.44 |
| Cabbage red | 39.2 | 28.4 | 47.7 | 35.7 | 1.72 | 1.34 | 1.93 | 1.53 |
| Cabbage savoy | 31.7 | 25.9 | 38.2 | 33.9 | 1.22 | 0.87 | 1.45 | 1.06 |
| Cauliflower | 32.5 | 27.2 | 40.7 | 34.6 | 1.35 | 0.99 | 1.58 | 1.18 |
| Collard | 34.2 | 27.1 | 41.1 | 33.5 | 1.37 | 1.01 | 1.61 | 1.25 |
| Kale Chinese | 29.5 | 24.2 | 35.3 | 31.6 | 1.08 | 0.72 | 1.31 | 0.91 |
| Kale red | 28.2 | 23.2 | 36.8 | 31.1 | 1.15 | 0.80 | 1.38 | 0.99 |
| Kale Tucsan | 38.6 | 33.8 | 49.5 | 41.4 | 1.79 | 1.43 | 2.02 | 1.62 |
| Kohlrabi purple | 37.2 | 31.2 | 46.2 | 38.6 | 1.62 | 1.27 | 1.82 | 1.46 |
| Cabbage Chinese | 33.9 | 26.4 | 39.4 | 32.7 | 1.28 | 0.93 | 1.51 | 1.12 |
| Komatsuna red | 31.5 | 28.2 | 44.2 | 36.6 | 1.52 | 1.17 | 1.75 | 1.36 |
| Mizuna | 38.4 | 30.7 | 44.8 | 36.3 | 1.55 | 1.20 | 1.78 | 1.39 |
| Pak choy | 35.2 | 29.1 | 40.3 | 36.1 | 1.33 | 0.97 | 1.56 | 1.65 |
| Rapini | 38.6 | 28.8 | 38.2 | 33.9 | 1.22 | 0.87 | 1.45 | 1.06 |
| Turnip | 41.8 | 31.4 | 43.3 | 36.2 | 1.45 | 1.12 | 1.71 | 1.31 |
| Mustard Dijon | 34.9 | 24.3 | 42.5 | 33.1 | 1.44 | 1.08 | 1.67 | 1.27 |
| Mustard red | 39.1 | 31.1 | 42.8 | 37.1 | 1.45 | 1.10 | 1.68 | 1.29 |
| Rutabaga | 31.7 | 23.5 | 38.7 | 29.7 | 1.25 | 0.89 | 1.48 | 1.08 |
| Tatsoi | 41.2 | 33.1 | 46.3 | 39.1 | 1.63 | 1.27 | 1.86 | 1.46 |
| LEDs | | | *** | | | | ** | |
| Varieties | ** | * | * | * | * | ** | *** | * |
| LEDs X Varieties | | | * | | | | ** | |

**Supplemental Table 3.** Growth parameters (Fresh weight (g), and Dry weight (%)) of of 21 varieties of Brassica microgreens represented 5 species grown under different light-emitting diodes (LEDs) ratio (%) of red:blue 80:20 ($R_{80}$:$B_{20}$), red:blue 20:80 ($R_{20}$:$B_{80}$), red:green:blue 70:10:20 ($R_{70}$:$G_{10}$:$B_{20}$), or red:green:blue 20:10:20 ($R_{20}$:$G_{10}$:$B_{70}$). Mean±SE values are based on a representative sample from each treatment across three experimental replications. NS, *, **, ***Nonsignificant or significant at P ≤ 0.05, 0.01, or 0.001, respectively.

| Commercial name | Fresh weight (g) | | | | Dry weight (%) | | | |
|---|---|---|---|---|---|---|---|---|
| | $R_{80}$:$B_{20}$ | $R_{20}$:$B_{80}$ | $R_{70}$:$G_{10}$:$B_{20}$ | $R_{20}$:$G_{10}$:$B_{70}$ | $R_{80}$:$B_{20}$ | $R_{20}$:$B_{80}$ | $R_{70}$:$G_{10}$:$B_{20}$ | $R_{20}$:$G_{10}$:$B_{70}$ |
| Broccoli | 0.38 | 0.30 | 0.42 | 0.36 | 5.96 | 5.32 | 6.55 | 5.65 |
| Brussel sprouts | 0.36 | 0.30 | 0.38 | 0.35 | 5.88 | 5.10 | 5.80 | 5.44 |
| Cabbage green | 0.40 | 0.33 | 0.44 | 0.39 | 6.2 | 5.46 | 6.25 | 5.79 |
| Cabbage red | 0.40 | 0.32 | 0.46 | 0.39 | 6.20 | 5.43 | 6.18 | 5.77 |
| Cabbage savoy | 0.34 | 0.28 | 0.36 | 0.33 | 5.67 | 5.01 | 6.17 | 5.34 |
| Cauliflower | 0.35 | 0.29 | 0.39 | 0.35 | 5.78 | 5.11 | 6.23 | 5.44 |
| Collard | 0.36 | 0.29 | 0.39 | 0.35 | 5.83 | 5.19 | 6.46 | 5.53 |
| Kale Chinese | 0.32 | 0.26 | 0.34 | 0.31 | 5.45 | 4.79 | 5.94 | 5.12 |
| Kale red | 0.31 | 0.25 | 0.35 | 0.31 | 5.38 | 4.74 | 5.99 | 5.07 |
| Kale Tucsan | 0.41 | 0.36 | 0.47 | 0.42 | 6.45 | 5.78 | 6.90 | 6.12 |
| Kohlrabi purple | 0.39 | 0.33 | 0.44 | 0.39 | 6.23 | 5.55 | 6.65 | 5.89 |
| Cabbage Chinese | 0.35 | 0.29 | 0.37 | 0.34 | 5.76 | 4.98 | 5.70 | 5.32 |
| Komatsuna red | 0.34 | 0.30 | 0.41 | 0.36 | 5.85 | 5.24 | 6.57 | 5.57 |
| Mizuna | 0.40 | 0.33 | 0.43 | 0.38 | 6.19 | 5.36 | 5.80 | 5.69 |
| Pak choy | 0.37 | 0.31 | 0.39 | 0.36 | 5.96 | 5.24 | 6.14 | 5.57 |
| Rapini | 0.39 | 0.31 | 0.38 | 0.35 | 6.01 | 5.31 | 6.33 | 5.65 |
| Turnip | 0.42 | 0.34 | 0.42 | 0.38 | 6.32 | 5.46 | 5.82 | 5.79 |
| Mustard Dijon | 0.36 | 0.28 | 0.41 | 0.35 | 5.80 | 5.10 | 6.13 | 5.44 |
| Mustard red | 0.40 | 0.33 | 0.42 | 0.38 | 6.22 | 5.37 | 5.78 | 5.70 |
| Rutabaga | 0.33 | 0.26 | 0.37 | 0.32 | 5.53 | 4.94 | 6.40 | 5.27 |
| Tatsoi | 0.42 | 0.35 | 0.45 | 0.41 | 6.45 | 5.67 | 6.34 | 6.00 |
| LEDs | | | ** | | | | ** | |
| Varieties | ** | *** | ** | ** | * | ** | * | * |
| LEDs X Varieties | | | ** | | | | ** | |

**Supplemental Table 4.** Mineral composition and content of macroelements; P and K (mg/100 g FW)) of of 21 varieties of Brassica microgreens represented 5 species grown under different light-emitting diodes (LEDs) ratio (%) of red:blue 80:20 ($R_{80}:B_{20}$), red:blue 20:80 ($R_{20}:B_{80}$), red:green:blue 70:10:20 ($R_{70}:G_{10}:B_{20}$), or red:green:blue 20:10:20 ($R_{20}:G_{10}:B_{70}$). Mean±SE values are based on a representative sample from each treatment across three experimental replications. NS, *, **, ***Nonsignificant or significant at P ≤ 0.05, 0.01, or 0.001, respectively.

| Commercial name | P (mg/100 g FW) | | | | K (mg/100 g FW) | | | |
|---|---|---|---|---|---|---|---|---|
| | $R_{80}:B_{20}$ | $R_{20}:B_{80}$ | $R_{70}:G_{10}:B_{20}$ | $R_{20}:G_{10}:B_{70}$ | $R_{80}:B_{20}$ | $R_{20}:B_{80}$ | $R_{70}:G_{10}:B_{20}$ | $R_{20}:G_{10}:B_{70}$ |
| Broccoli | 57 | 65 | 59 | 60 | 300 | 371 | 319 | 335 |
| Brussel sprouts | 53 | 68 | 57 | 60 | 287 | 365 | 308 | 325 |
| Cabbage green | 61 | 69 | 62 | 64 | 183 | 199 | 183 | 188 |
| Cabbage red | 59 | 71 | 62 | 64 | 218 | 254 | 224 | 233 |
| Cabbage savoy | 56 | 68 | 59 | 61 | 206 | 236 | 210 | 218 |
| Cauliflower | 56 | 72 | 60 | 63 | 199 | 272 | 218 | 234 |
| Collard | 66 | 76 | 67 | 70 | 274 | 300 | 275 | 283 |
| Kale Chinese | 62 | 69 | 64 | 66 | 220 | 298 | 244 | 261 |
| Kale red | 50 | 65 | 55 | 58 | 317 | 383 | 336 | 350 |
| Kale Tucsan | 60 | 69 | 63 | 65 | 268 | 310 | 280 | 289 |
| Kohlrabi purple | 65 | 75 | 68 | 70 | 306 | 362 | 322 | 334 |
| Cabbage Chinese | 63 | 72 | 65 | 67 | 211 | 280 | 231 | 245 |
| Komatsuna red | 62 | 77 | 66 | 70 | 298 | 370 | 320 | 335 |
| Mizuna | 52 | 71 | 58 | 62 | 315 | 402 | 341 | 359 |
| Pak choy | 51 | 60 | 54 | 56 | 282 | 290 | 284 | 286 |
| Rapini | 68 | 85 | 73 | 77 | 324 | 411 | 350 | 368 |
| Turnip | 49 | 68 | 55 | 59 | 296 | 301 | 297 | 299 |
| Mustard Dijon | 57 | 68 | 60 | 62 | 323 | 400 | 346 | 362 |
| Mustard red | 43 | 49 | 44 | 46 | 257 | 311 | 273 | 284 |
| Rutabaga | 58 | 75 | 63 | 67 | 206 | 249 | 218 | 228 |
| Tatsoi | 62 | 71 | 64 | 66 | 275 | 359 | 300 | 318 |
| LEDs | | | * | | | | ** | |
| Varieties | ** | * | ** | ** | * | ** | * | * |
| LEDs X Varieties | | | * | | | | ** | |

**Supplemental Table 5.** Mineral composition and content of macroelements; Ca and Mg (mg/100 g FW)) of 21 varieties of Brassica microgreens represented 5 species grown under different light-emitting diodes (LEDs) ratio (%) of red:blue 80:20 ($R_{80}:B_{20}$), red:blue 20:80 ($R_{20}:B_{80}$), red:green:blue 70:10:20 ($R_{70}:G_{10}:B_{20}$), or red:green:blue 20:10:20 ($R_{20}:G_{10}:B_{70}$). Mean±SE values are based on a representative sample from each treatment across three experimental replications. NS, *, **, ***Nonsignificant or significant at P ≤ 0.05, 0.01, or 0.001, respectively.

| Commercial name | Ca (mg/100 g FW) | | | | Na (mg/100 g FW) | | | |
|---|---|---|---|---|---|---|---|---|
| | $R_{80}:B_{20}$ | $R_{20}:B_{80}$ | $R_{70}:G_{10}:B_{20}$ | $R_{20}:G_{10}:B_{70}$ | $R_{80}:B_{20}$ | $R_{20}:B_{80}$ | $R_{70}:G_{10}:B_{20}$ | $R_{20}:G_{10}:B_{70}$ |
| Broccoli | 84 | 112 | 92 | 98 | 50 | 59 | 50 | 53 |
| Brussel sprouts | 62 | 75 | 65 | 68 | 54 | 55 | 51 | 52 |
| Cabbage green | 84 | 100 | 87 | 91 | 74 | 79 | 69 | 72 |
| Cabbage red | 79 | 100 | 84 | 89 | 42 | 46 | 40 | 42 |
| Cabbage savoy | 85 | 105 | 89 | 94 | 57 | 68 | 56 | 59 |
| Cauliflower | 79 | 100 | 84 | 89 | 76 | 83 | 72 | 75 |
| Collard | 62 | 74 | 64 | 67 | 47 | 49 | 44 | 45 |
| Kale Chinese | 39 | 52 | 43 | 46 | 39 | 45 | 40 | 41 |
| Kale red | 42 | 53 | 45 | 48 | 40 | 52 | 43 | 45 |
| Kale Tucsan | 54 | 58 | 55 | 56 | 46 | 51 | 46 | 48 |
| Kohlrabi purple | 63 | 72 | 65 | 67 | 45 | 53 | 46 | 48 |
| Cabbage Chinese | 69 | 78 | 71 | 73 | 25 | 27 | 24 | 25 |
| Komatsuna red | 48 | 63 | 53 | 56 | 24 | 38 | 28 | 31 |
| Mizuna | 36 | 54 | 42 | 45 | 39 | 46 | 40 | 42 |
| Pak choy | 45 | 57 | 49 | 51 | 28 | 35 | 31 | 32 |
| Rapini | 85 | 94 | 87 | 89 | 58 | 65 | 58 | 60 |
| Turnip | 42 | 52 | 45 | 47 | 18 | 29 | 22 | 24 |
| Mustard Dijon | 40 | 50 | 43 | 45 | 30 | 31 | 29 | 30 |
| Mustard red | 47 | 60 | 51 | 54 | 28 | 34 | 29 | 31 |
| Rutabaga | 53 | 61 | 55 | 57 | 43 | 57 | 46 | 49 |
| Tatsoi | 43 | 48 | 44 | 45 | 35 | 39 | 35 | 36 |
| LEDs | | | ** | | | | ** | |
| Varieties | * | ** | * | * | * | * | * | ** |
| LEDs X Varieties | | | * | | | | ** | |

**Supplemental Table 6.** Mineral composition and content of microelements; Fe and Zn (mg/100 g FW)) of of 21 varieties of Brassica microgreens represented 5 species grown under different light-emitting diodes (LEDs) ratio (%) of red:blue 80:20 ($R_{80}$:$B_{20}$), red:blue 20:80 ($R_{20}$:$B_{80}$), red:green:blue 70:10:20 ($R_{70}$:$G_{10}$:$B_{20}$), or red:green:blue 20:10:20 ($R_{20}$:$G_{10}$:$B_{70}$). Mean±SE values are based on a representative sample from each treatment across three experimental replications. NS, *, **, ***Nonsignificant or significant at P ≤ 0.05, 0.01, or 0.001, respectively.

| Commercial name | Fe (mg/100 g FW) | | | | Zn (mg/100 g FW) | | | |
|---|---|---|---|---|---|---|---|---|
| | $R_{80}$:$B_{20}$ | $R_{20}$:$B_{80}$ | $R_{70}$:$G_{10}$:$B_{20}$ | $R_{20}$:$G_{10}$:$B_{70}$ | $R_{80}$:$B_{20}$ | $R_{20}$:$B_{80}$ | $R_{70}$:$G_{10}$:$B_{20}$ | $R_{20}$:$G_{10}$:$B_{70}$ |
| Broccoli | 0.73 | 0.85 | 0.47 | 0.47 | 0.47 | 0.47 | 0.47 | 0.47 |
| Brussel sprouts | 0.61 | 0.72 | 0.36 | 0.36 | 0.36 | 0.36 | 0.36 | 0.36 |
| Cabbage green | 0.66 | 0.77 | 0.36 | 0.36 | 0.36 | 0.36 | 0.36 | 0.36 |
| Cabbage red | 0.68 | 0.79 | 0.44 | 0.44 | 0.44 | 0.44 | 0.44 | 0.44 |
| Cabbage savoy | 0.59 | 0.68 | 0.35 | 0.35 | 0.35 | 0.35 | 0.35 | 0.35 |
| Cauliflower | 0.66 | 0.76 | 0.37 | 0.37 | 0.37 | 0.37 | 0.37 | 0.37 |
| Collard | 0.70 | 0.80 | 0.48 | 0.48 | 0.48 | 0.48 | 0.48 | 0.48 |
| Kale Chinese | 0.70 | 0.80 | 0.41 | 0.41 | 0.41 | 0.41 | 0.41 | 0.41 |
| Kale red | 0.51 | 0.59 | 0.32 | 0.32 | 0.32 | 0.32 | 0.32 | 0.32 |
| Kale Tucsan | 0.77 | 0.88 | 0.42 | 0.42 | 0.42 | 0.42 | 0.42 | 0.42 |
| Kohlrabi purple | 0.78 | 0.90 | 0.48 | 0.48 | 0.48 | 0.48 | 0.48 | 0.48 |
| Cabbage Chinese | 0.70 | 0.81 | 0.44 | 0.44 | 0.44 | 0.44 | 0.44 | 0.44 |
| Komatsuna red | 0.78 | 0.90 | 0.43 | 0.43 | 0.43 | 0.43 | 0.43 | 0.43 |
| Mizuna | 0.61 | 0.71 | 0.32 | 0.32 | 0.32 | 0.32 | 0.32 | 0.32 |
| Pak choy | 0.54 | 0.68 | 0.44 | 0.44 | 0.44 | 0.44 | 0.44 | 0.44 |
| Rapini | 0.88 | 1.05 | 0.68 | 0.68 | 0.68 | 0.68 | 0.68 | 0.68 |
| Turnip | 0.62 | 0.73 | 0.40 | 0.40 | 0.40 | 0.40 | 0.40 | 0.40 |
| Mustard Dijon | 0.60 | 0.70 | 0.32 | 0.32 | 0.32 | 0.32 | 0.32 | 0.32 |
| Mustard red | 0.68 | 0.84 | 0.27 | 0.27 | 0.27 | 0.27 | 0.27 | 0.27 |
| Rutabaga | 0.58 | 0.72 | 0.36 | 0.36 | 0.36 | 0.36 | 0.36 | 0.36 |
| Tatsoi | 0.61 | 0.75 | 0.44 | 0.44 | 0.44 | 0.44 | 0.44 | 0.44 |
| LEDs | *** | | | | ** | | | |
| Varieties | * | *** | * | ** | * | ** | *** | * |
| LEDs X Varieties | ** | | | | ** | | | |

**Supplemental Table 7.** Mineral composition and content of microelements; Cu and Mn (mg/100 g FW)) of 21 varieties of Brassica microgreens represented 5 species grown under different light-emitting diodes (LEDs) ratio (%) of red:blue 80:20 ($R_{80}$:$B_{20}$), red:blue 20:80 ($R_{20}$:$B_{80}$), red:green:blue 70:10:20 ($R_{70}$:$G_{10}$:$B_{20}$), or red:green:blue 20:10:20 ($R_{20}$:$G_{10}$:$B_{70}$). Mean±SE values are based on a representative sample from each treatment across three experimental replications. NS, *, **, ***Nonsignificant or significant at P ≤ 0.05, 0.01, or 0.001, respectively.

| Commercial name | Cu (mg/100 g FW) | | | | Mn (mg/100 g FW) | | | |
|---|---|---|---|---|---|---|---|---|
| | $R_{80}$:$B_{20}$ | $R_{20}$:$B_{80}$ | $R_{70}$:$G_{10}$:$B_{20}$ | $R_{20}$:$G_{10}$:$B_{70}$ | $R_{80}$:$B_{20}$ | $R_{20}$:$B_{80}$ | $R_{70}$:$G_{10}$:$B_{20}$ | $R_{20}$:$G_{10}$:$B_{70}$ |
| Broccoli | 0.10 | 0.16 | 0.11 | 0.13 | 0.40 | 0.49 | 0.41 | 0.46 |
| Brussel sprouts | 0.09 | 0.15 | 0.10 | 0.12 | 0.41 | 0.50 | 0.42 | 0.46 |
| Cabbage green | 0.05 | 0.11 | 0.06 | 0.08 | 0.35 | 0.43 | 0.37 | 0.41 |
| Cabbage red | 0.09 | 0.15 | 0.10 | 0.12 | 0.33 | 0.42 | 0.35 | 0.39 |
| Cabbage savoy | 0.06 | 0.09 | 0.08 | 0.09 | 0.42 | 0.49 | 0.44 | 0.48 |
| Cauliflower | 0.07 | 0.10 | 0.09 | 0.09 | 0.34 | 0.41 | 0.36 | 0.39 |
| Collard | 0.10 | 0.14 | 0.12 | 0.13 | 0.39 | 0.46 | 0.41 | 0.44 |
| Kale Chinese | 0.06 | 0.10 | 0.04 | 0.05 | 0.30 | 0.36 | 0.27 | 0.30 |
| Kale red | 0.06 | 0.10 | 0.04 | 0.05 | 0.35 | 0.42 | 0.32 | 0.36 |
| Kale Tucsan | 0.09 | 0.15 | 0.07 | 0.09 | 0.48 | 0.57 | 0.46 | 0.50 |
| Kohlrabi purple | 0.12 | 0.17 | 0.08 | 0.11 | 0.42 | 0.50 | 0.39 | 0.43 |
| Cabbage Chinese | 0.04 | 0.10 | 0.05 | 0.07 | 0.33 | 0.41 | 0.34 | 0.38 |
| Komatsuna red | 0.08 | 0.14 | 0.05 | 0.07 | 0.37 | 0.45 | 0.34 | 0.37 |
| Mizuna | 0.06 | 0.11 | 0.03 | 0.05 | 0.39 | 0.47 | 0.36 | 0.39 |
| Pak choy | 0.06 | 0.16 | 0.13 | 0.15 | 0.32 | 0.44 | 0.40 | 0.41 |
| Rapini | 0.14 | 0.23 | 0.23 | 0.24 | 0.45 | 0.57 | 0.54 | 0.58 |
| Turnip | 0.08 | 0.13 | 0.09 | 0.10 | 0.44 | 0.53 | 0.45 | 0.50 |
| Mustard Dijon | 0.07 | 0.12 | 0.03 | 0.05 | 0.43 | 0.51 | 0.39 | 0.43 |
| Mustard red | 0.06 | 0.16 | 0.05 | 0.08 | 0.25 | 0.36 | 0.24 | 0.28 |
| Rutabaga | 0.08 | 0.17 | 0.13 | 0.14 | 0.43 | 0.56 | 0.49 | 0.53 |
| Tatsoi | 0.05 | 0.14 | 0.08 | 0.10 | 0.32 | 0.43 | 0.35 | 0.39 |
| LEDs | | | *** | | ** | | | |
| Varieties | * | ** | ** | * | ** | * | | ** |
| LEDs X Varieties | | | * | | | | *** | |

**Supplemental Table 8.** Vitamins concentrations (Phylloquinone (ug/ g FW) and α-tocopherol (mg/100 g FW)) of 21 varieties of Brassica microgreens represented 5 species grown under different light-emitting diodes (LEDs) ratio (%) of red:blue 80:20 ($R_{80}$:$B_{20}$), red:blue 20:80 ($R_{20}$:$B_{80}$), red:green:blue 70:10:20 ($R_{70}$:$G_{10}$:$B_{20}$), or red:green:blue 20:10:20 ($R_{20}$:$G_{10}$:$B_{70}$). Mean±SE values are based on a representative sample from each treatment across three experimental replications. NS, *, **, ***Nonsignificant or significant at P ≤ 0.05, 0.01, or 0.001, respectively.

| Commercial name | Phylloquinone (ug/ g FW) | | | | α-tocopherol (mg/100 g FW) | | | |
|---|---|---|---|---|---|---|---|---|
| | $R_{80}$:$B_{20}$ | $R_{20}$:$B_{80}$ | $R_{70}$:$G_{10}$:$B_{20}$ | $R_{20}$:$G_{10}$:$B_{70}$ | $R_{80}$:$B_{20}$ | $R_{20}$:$B_{80}$ | $R_{70}$:$G_{10}$:$B_{20}$ | $R_{20}$:$G_{10}$:$B_{70}$ |
| Broccoli | 1.9 | 2.3 | 2.0 | 2.2 | 16.2 | 20.0 | 17.3 | 19.0 |
| Brussel sprouts | 1.3 | 2.0 | 1.5 | 1.6 | 16.3 | 20.2 | 17.5 | 19.2 |
| Cabbage green | 1.1 | 1.9 | 1.4 | 1.4 | 13.4 | 16.6 | 14.3 | 15.8 |
| Cabbage red | 2.2 | 2.5 | 2.3 | 2.6 | 27.6 | 34.0 | 29.5 | 32.4 |
| Cabbage savoy | 2.3 | 2.8 | 2.5 | 2.7 | 19.5 | 24.1 | 20.9 | 23.0 |
| Cauliflower | 1.7 | 2.1 | 1.8 | 2.0 | 14.9 | 18.4 | 16.0 | 17.6 |
| Collard | 1.9 | 2.3 | 2.0 | 2.0 | 17.5 | 21.6 | 18.5 | 19.0 |
| Kale Chinese | 1.2 | 1.5 | 1.3 | 1.3 | 13.5 | 16.7 | 14.3 | 14.7 |
| Kale red | 1.4 | 1.7 | 1.5 | 1.5 | 16.4 | 20.3 | 17.4 | 17.9 |
| Kale Tucsan | 1.6 | 2.0 | 1.7 | 1.7 | 18.4 | 22.7 | 19.4 | 20.0 |
| Kohlrabi purple | 2.5 | 3.1 | 2.6 | 2.7 | 16.7 | 20.6 | 17.6 | 18.1 |
| Cabbage Chinese | 1.6 | 2.1 | 1.8 | 1.9 | 17.8 | 21.9 | 19.0 | 20.9 |
| Komatsuna red | 2.1 | 2.6 | 2.2 | 2.3 | 23.7 | 29.2 | 25.0 | 25.7 |
| Mizuna | 1.3 | 1.6 | 1.4 | 1.4 | 25.1 | 31.0 | 26.5 | 27.3 |
| Pak choy | 1.7 | 2.1 | 1.8 | 1.9 | 17.2 | 21.3 | 18.8 | 20.0 |
| Rapini | 1.1 | 1.4 | 1.2 | 1.4 | 17.6 | 21.7 | 18.3 | 20.9 |
| Turnip | 1.4 | 1.7 | 1.6 | 1.6 | 17.5 | 21.6 | 19.7 | 20.2 |
| Mustard Dijon | 1.3 | 1.6 | 1.4 | 1.5 | 20.2 | 25.0 | 22.0 | 23.5 |
| Mustard red | 2.0 | 2.5 | 2.2 | 2.4 | 23.3 | 28.8 | 25.4 | 27.1 |
| Rutabaga | 1.6 | 2.0 | 1.7 | 1.9 | 31.6 | 39.0 | 32.9 | 37.6 |
| Tatsoi | 1.3 | 1.6 | 1.5 | 1.5 | 26.1 | 32.2 | 29.3 | 30.1 |
| LEDs | | | *** | | | | ** | |
| Varieties | *** | * | ** | * | *** | * | ** | ** |
| LEDs X Varieties | | | * | | | | ** | |

**Supplemental Table 9.** Vitamin and carotenoid concentrations (Total Ascorbic Acid (mg/100 g FW) and β-carotene (mg/100 g FW)) of 21 varieties of Brassica microgreens represented 5 species grown under different light-emitting diodes (LEDs) ratio (%) of red:blue 80:20 ($R_{80}$:$B_{20}$), red:blue 20:80 ($R_{20}$:$B_{80}$), red:green:blue 70:10:20 ($R_{70}$:$G_{10}$:$B_{20}$), or red:green:blue 20:10:20 ($R_{20}$:$G_{10}$:$B_{70}$). Mean±SE values are based on a representative sample from each treatment across three experimental replications. NS, *, **, ***Nonsignificant or significant at P ≤ 0.05, 0.01, or 0.001, respectively.

| Commercial name | Total Ascorbic Acid (mg/100 g FW) | | | | β-carotene (mg/100 g FW) | | | |
|---|---|---|---|---|---|---|---|---|
| | $R_{80}$:$B_{20}$ | $R_{20}$:$B_{80}$ | $R_{70}$:$G_{10}$:$B_{20}$ | $R_{20}$:$G_{10}$:$B_{70}$ | $R_{80}$:$B_{20}$ | $R_{20}$:$B_{80}$ | $R_{70}$:$G_{10}$:$B_{20}$ | $R_{20}$:$G_{10}$:$B_{70}$ |
| Broccoli | 79.2 | 97.7 | 84.6 | 93.2 | 6.4 | 8.0 | 6.9 | 7.6 |
| Brussel sprouts | 89.7 | 110.7 | 95.8 | 105.6 | 5.1 | 6.3 | 5.5 | 6.0 |
| Cabbage green | 111.3 | 137.4 | 118.9 | 131.0 | 9.0 | 11.1 | 9.6 | 10.6 |
| Cabbage red | 119.3 | 147.2 | 127.4 | 140.4 | 9.3 | 11.4 | 9.9 | 10.9 |
| Cabbage savoy | 116.6 | 143.9 | 124.6 | 137.3 | 10.5 | 13.0 | 11.2 | 12.4 |
| Cauliflower | 90.1 | 111.2 | 96.3 | 106.1 | 7.4 | 9.2 | 7.9 | 8.7 |
| Collard | 70.7 | 87.2 | 74.6 | 76.8 | 4.9 | 6.1 | 5.3 | 5.4 |
| Kale Chinese | 73.2 | 90.4 | 77.4 | 79.6 | 7.3 | 9.0 | 7.7 | 8.0 |
| Kale red | 59.1 | 72.9 | 62.4 | 64.2 | 6.1 | 7.5 | 6.4 | 6.6 |
| Kale Tucsan | 69.4 | 85.6 | 73.2 | 75.4 | 5.1 | 6.3 | 5.4 | 5.6 |
| Kohlrabi purple | 73.0 | 90.1 | 77.1 | 79.3 | 6.3 | 7.7 | 6.6 | 6.8 |
| Cabbage Chinese | 110.4 | 136.3 | 118.0 | 130.0 | 8.3 | 10.2 | 8.9 | 9.8 |
| Komatsuna red | 92.3 | 113.9 | 97.5 | 100.3 | 6.7 | 8.3 | 7.1 | 7.3 |
| Mizuna | 44.4 | 54.8 | 46.9 | 48.3 | 6.0 | 7.4 | 6.3 | 6.5 |
| Pak choy | 40.6 | 50.0 | 44.2 | 47.0 | 6.4 | 8.0 | 7.0 | 7.5 |
| Rapini | 79.1 | 97.6 | 82.2 | 94.0 | 8.3 | 10.2 | 8.6 | 9.9 |
| Turnip | 61.7 | 76.1 | 69.4 | 71.2 | 8.1 | 10.0 | 9.2 | 9.4 |
| Mustard Dijon | 78.5 | 96.9± | 85.5 | 91.1 | 5.5 | 6.8 | 6.0 | 6.4 |
| Mustard red | 48.9 | 60.4 | 53.3 | 56.8 | 3.8 | 4.7 | 4.1 | 4.4 |
| Rutabaga | 78.0 | 96.3 | 81.1 | 92.7 | 6.0 | 7.4 | 6.2 | 7.1 |
| Tatsoi | 88.4 | 109.0 | 99.5 | 102.0 | 9.5 | 11.7 | 10.6 | 10.9 |
| LEDs | | | *** | | | | ** | |
| Varieties | * | * | * | ** | * | ** | * | ** |
| LEDs X Varieties | | | * | | | | ** | |
